# A positive feedback loop controls *Toxoplasma* chronic differentiation

**DOI:** 10.1101/2022.04.06.487076

**Authors:** M. Haley Licon, Christopher J. Giuliano, Sundeep Chakladar, Lindsey Shallberg, Benjamin S. Waldman, Christopher A. Hunter, Sebastian Lourido

## Abstract

Successful infection strategies must balance pathogen amplification and persistence. In *Toxoplasma gondii*, this is accomplished through differentiation into dedicated cyst-forming chronic stages that avoid clearance by the host immune system. The transcription factor BFD1 is both necessary and sufficient for stage conversion; however, its regulation is not understood. We examine five factors transcriptionally activated by BFD1. One of these is a cytosolic RNA-binding protein of the CCCH-type zinc finger family, which we name BFD2. Parasites lacking BFD2 fail to induce BFD1 and are consequently unable to fully differentiate in culture or in mice. BFD2 interacts with the *BFD1* transcript in a stress-dependent manner. Deletion of BFD2 reduces BFD1 protein levels, but not mRNA abundance. The reciprocal effects on BFD2 transcription and BFD1 translation outline a positive feedback loop that enforces commitment to differentiation. BFD2 helps explain how parasites commit to the chronic gene-expression program and elucidates how the balance between proliferation and persistence is achieved over the course of infection.

## INTRODUCTION

The ability to establish productive infection requires a trade-off between pathogen replication and persistence. While replication increases the likelihood of transmission, the resultant tissue damage can drive host immunity and promote pathogen clearance. At the same time, uncontrolled pathogen proliferation may be fatal for the host and altogether eliminate the reservoir of infection (Cressler et al., 2016). To balance these outcomes, many protozoan parasites have developed slow-growing chronic stages that ensure parasite survival during unfavorable growth conditions and frequently exhibit reduced immunogenicity (Barrett et al., 2019). The equilibrium between proliferation and latency thereby ensures host survival and parasite transmission.

For apicomplexans of the *Sarcocystidae* family, persistence is achieved by forming large intracellular structures (tissue cysts) within the host that contain dedicated chronic stages known as bradyzoites. Whereas infected animals can survive long-term harboring these cysts, active replication of the proliferative forms (tachyzoites) results in severe, potentially deadly disease. The most widespread of these infections is caused by *Toxoplasma gondii*, which chronically infects up to one quarter of humans on Earth (Montoya and Liesenfeld, 2004). During acute infection, *T. gondii* tachyzoites disseminate throughout the body and cause pathology through host-cell lysis. A proportion of tachyzoites give rise to bradyzoites, which encyst in brain and muscle tissue (Dubey et al., 1998). Most infections are controlled by the host immune response; however, reactivation of latent cysts can occur during periods of weakened immune function, such as during immunosuppressive therapy or in patients with advanced HIV infection (Derouin et al., 2008; Grant et al., 1990; Jones et al., 2015; Luft and Remington, 1992). Additionally, none of the clinical therapies target chronic stages (Alday and Doggett, 2017), making it currently impossible to eradicate *Toxoplasma* from the host.

While the physiological triggers of differentiation remain poorly understood, in cell culture the transition can be induced by a variety of exogenous stressors, including alkaline pH, heat shock, small molecules, and nutrient restriction (Skariah et al., 2010; Soête et al., 1994). Early in the transition, the parasitophorous vacuole is remodeled into a glycan-rich cyst wall, containing many stage-specific proteins of unknown function (Ferguson and Hutchison, 1987; Tu et al., 2019, 2020). Bradyzoite metabolism shifts from aerobic respiration to anaerobic glycolysis and parasites accumulate cytoplasmic starch granules (Denton et al., 1996; Fox et al., 2004; Shukla et al., 2018). These changes coincide with a global restructuring of *Toxoplasma* gene expression (Behnke et al., 2008; Buchholz et al., 2011; Cleary et al., 2002; Pittman et al., 2014; Radke et al., 2005; Waldman et al., 2020), indicating that transcriptional networks likely play a major role in stage conversion. Accordingly, examples of chromatin-binding (Bougdour et al., 2009; Farhat et al., 2020; Maubon et al., 2010; Saksouk et al., 2005), RNA-binding (Gissot et al., 2013, 2017; Liu et al., 2014; Vanchinathan et al., 2005), and DNA-binding (Hong et al., 2017; Huang et al., 2017; Radke et al., 2013, 2018; Walker et al., 2013) proteins have all been identified with both activating and antagonistic effects. Individual gene knockouts impacted overall rates of cyst formation (Hong et al., 2017; Huang et al., 2017; Radke et al., 2013; Walker et al., 2013); however, it wasn’t until the discovery of BFD1 that chronic differentiation could be shown to depend on the activity of a single factor.

Bradyzoite Formation Deficient 1 (BFD1), a Myb-like transcription factor, was discovered by screening ∼200 *T. gondii* genes with putative nucleic acid-binding domains (Waldman et al., 2020). Despite being transcribed throughout the *Toxoplasma* asexual cycle, BFD1 protein is not detected in unstressed parasites, implying that it is translationally controlled. Parasites lacking BFD1 grow normally as tachyzoites, but enter a non-differentiated G1-arrested state under alkaline stress. Conversely, conditional BFD1 expression is sufficient for differentiation in the absence of stress and closely recapitulates the transcriptional changes observed in naturally-derived bradyzoites. The genome-wide BFD1 binding profile also reveals a preferential association with the promoters of many stage-specific genes, consistent with its function as a positive driver of the chronic gene-expression program.

While master regulators can be sufficient to induce developmental programs, they generally rely on the combined activities of other effectors to rewire existing cell states (Davis and Rebay, 2017). Inherent to these networks is a high degree of complexity involving hierarchical transcription, feedback circuits, and cooperative regulation. It is therefore unclear how other chronic-stage regulators intersect with BFD1. We previously showed that a number of genes containing putative RNA- and DNA-binding domains are targets of BFD1, suggesting a cascade of secondary differentiation factors. In the present study, we set out to unravel the regulatory network extending from BFD1 by characterizing these effectors. In doing so, we uncover a positive feedback loop that reinforces commitment to chronic differentiation.

## RESULTS

### Analysis of BFD1-regulated gene-expression factors

Analysis of previous data sets revealed five putative nucleic acid-binding proteins that are both upregulated during differentiation and directly bound in their promoters by BFD1, making them likely candidates for secondary effectors of the bradyzoite program (**Fig. 1A**). Two of these—*TGME49_306620* (*AP2IX-9*) and *TGME49_208020* (*AP2IB-1*)*—*are members of a well-characterized family of apicomplexan transcription factors, known as ApiAP2. *TGME49_311100, TGME49_224630*, and *TGME49_253790* encode CCCH-type zinc finger domains, commonly associated with RNA-binding. Whereas AP2IX-9 was described previously as an early-bradyzoite transcription factor (Hong et al., 2017; Radke et al., 2013), the other four genes were functionally uncharacterized. We therefore generated conditional knockdown strains for each candidate to assess its role in differentiation. In a parental line expressing the heterologous TIR1 auxin receptor, we endogenously tagged each candidate with mNeonGreen (mNG) and the minimal auxin-inducible degron (mAID), allowing both detection and regulated depletion of the gene product upon addition of indole-3-acetic acid (IAA) (Brown et al., 2018; Smith et al., 2021) (**Fig. 1B**). BFD1 was included as a positive control for a gene whose knockdown has a profound effect on differentiation.

**Figure 1.**
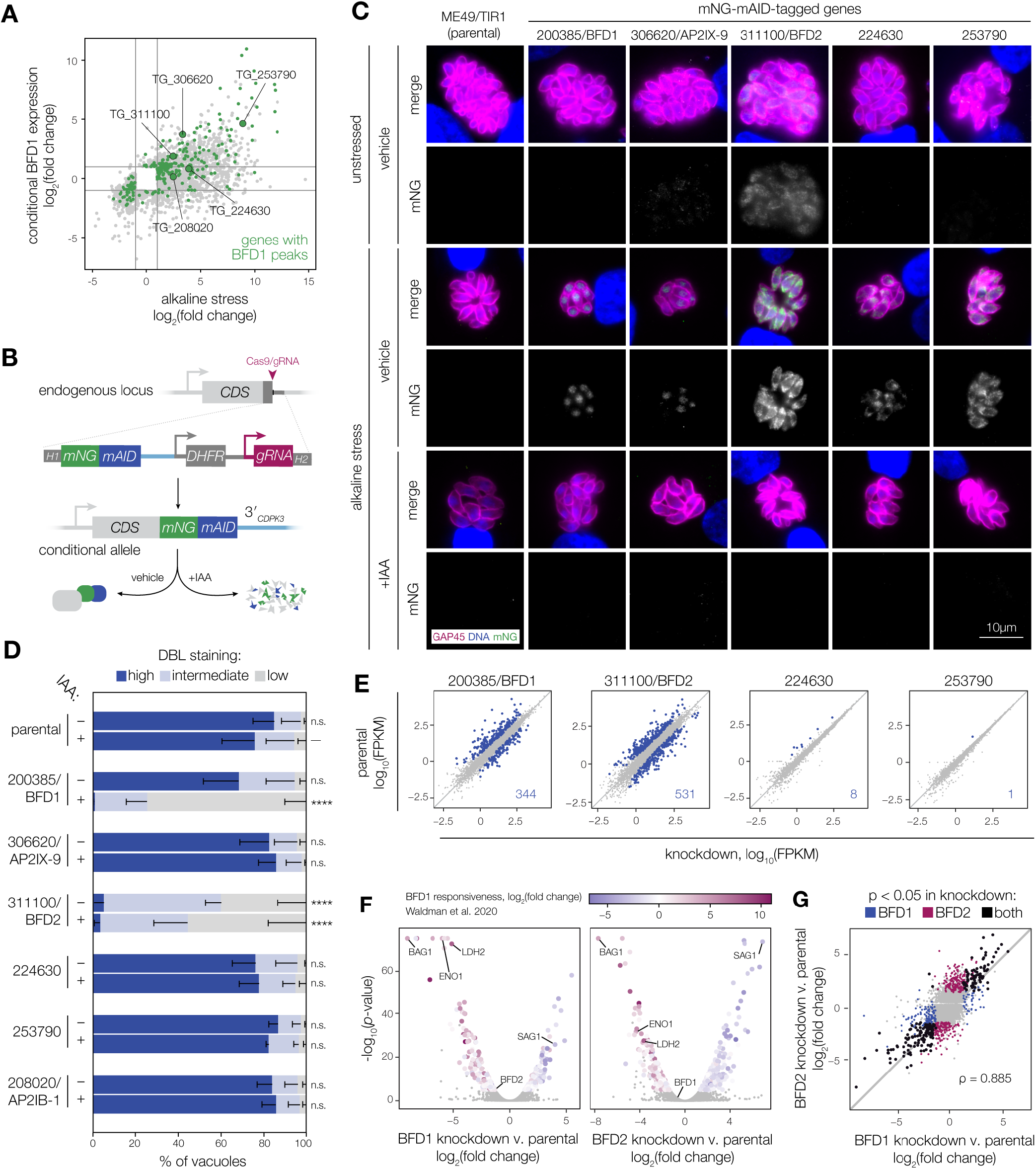
Screening for downstream effectors of BFD1. (**A**) Comparison of genes that are significantly regulated after 48 h under alkaline stress or conditional BFD1 expression (adjusted p < 0.001, 2-fold change or greater). Targets of BFD1 identified by CUT&RUN chromatin profiling are highlighted green (Waldman et al., 2020), including five genes that encode predicted RNA-or DNA-binding domains. (**B**) Generation of conditional knockdown strains. C-terminal tagging constructs were integrated downstream of each coding sequence (CDS) by Cas9-mediated homologous recombination and selected based on pyrimethamine resistance. Genes tagged with *-mNG-mAID* are targeted for proteasomal degradation upon treatment with IAA. Additional strain details are provided in Materials & Methods and **Fig. S4**. (**C**) Knockdown strains after 48 h under alkaline stress or unstressed conditions, treated with IAA or vehicle. GAP45 labels the parasite inner membrane complex (magenta) and DNA is stained with Hoechst (blue). Tagged genes are immunostained for mNG (green). The mNG channel was normalized internally for all samples from a given strain, using the parental as a negative control. For clarity, this channel is shown in greyscale below the corresponding merged image. (**D**) Differentiation of knockdown strains after 48 h of alkaline stress. Differentiation status was assessed by immunofluorescence using FITC-conjugated *Dolichos biflorus* lectin (DBL), which stains the bradyzoite cyst wall. Mean ± s.d. plotted for *n* = 3 biological replicates, with a minimum of 175 vacuoles counted per sample. One-way ANOVA performed with Dunnett’s method for multiple comparisons on percent of vacuoles scoring “DBL-high”, with each sample compared to the IAA-treated parental. \*\*\*\**p* < 0.0001 (**E–G**) Effects of candidate knockdown on the chronic-stage transcriptome. Data reflect changes after 96 h of alkaline stress and IAA, and are based on *n* = 2 independent infections. (**E**) Genes significantly affected by depletion of the indicated factors (adjusted *p*-value<0.05 calculated by DESeq2). The number of genes meeting the statistical threshold is indicated on each plot. Complete lists of genes provided in **Table S1. (F)** Genes significantly affected by BFD1 or BFD2 knockdown are colored by their log2(fold change) during conditional BFD1 expression (Waldman et al., 2020). (**G**) Comparison of the transcriptional effects of BFD1 and BFD2 depletion. Genes meeting the significance threshold in either sample are highlighted separately, with those reaching significance in both colored black. Spearman correlation was performed on the union of statistically regulated genes in both samples.

We first examined stress-induced expression and localization of each factor. Following 48 h of alkaline stress, we observed staining for BFD1 and four of the five candidates (**Fig. 1C**), with AP2IB-1 remaining below the limit of detection (not pictured). TGME49_311100 was uniquely expressed in acute stages, but clearly upregulated in response to stress. The three predicted RNA-binding proteins localized to the cytosol, whereas BFD1 and AP2IX-9 localized to the nuclei of stressed parasites, consistent with both their proposed functions as transcription factors and previous characterization (Radke et al., 2013; Waldman et al., 2020). All four detectable factors were visibly depleted following treatment with IAA (**Fig. 1C**).

To examine the impact of down-regulating each factor on stress-induced differentiation, parasites were allowed to invade host cells for 4 h under standard conditions, then switched to alkaline-stress media containing IAA or vehicle. After 48 h, vacuoles were labeled with *Dolichos biflorus* lectin (DBL), which binds N-acetylgalactosamine in the bradyzoite cyst wall. Differentiation was blindly scored based on DBL staining intensity using an automated image analysis pipeline (**Fig. 1D**). As expected, stage conversion was completely inhibited by depletion of BFD1 (Waldman et al., 2020). AP2IX-9—previously implicated as an antagonist of the chronic stage—showed no impact on stress-induced differentiation, consistent with published results (Hong et al., 2017; Radke et al., 2013). By contrast, parasites expressing the conditional allele of *TGME49_311100* showed a profound decrease in DBL-positive vacuoles under both vehicle and IAA treatment. Given the insensitivity of this effect to IAA, we conclude the activity of TGME49_311100 is likely disrupted by C-terminal tagging. Furthermore, our results implicate TGME49_311100 in the process of differentiation, leading us to name it Bradyzoite Formation Deficient 2 (BFD2).

### Transcriptional profiling reveals a common signature for BFD1 and BFD2 downregulation

Since DBL staining only describes one aspect of chronic differentiation, we subjected all of the mutants to RNA sequencing following 96 h of alkaline stress. Somewhat unexpectedly, few of the mutants exhibited any obvious signature that would indicate a role in transcriptional reprogramming (**Fig. 1E**). AP2IX-9 and AP2IB-1 in particular had no measurable effects when depleted from chronic stages (**Fig. S1**), despite clear evidence for the former in early-stage bradyzoite development (Hong et al., 2017; Radke et al., 2013). Additionally, although we did identify a small regulon for zinc finger protein TGME49_224630 (**Table S1**), the affected transcripts were largely limited to genes previously shown to be induced under alkaline stress but not responsive to BFD1 (Waldman et al., 2020). Thus, this likely represents a transcriptional response to alkaline stress rather than a bona fide component of the core bradyzoite program. By contrast, BFD1 knockdown had the expected impact on the chronic-stage transcriptome, which inversely mirrored the effects of conditionally upregulating BFD1 under standard conditions (**Fig. 1F**). Disrupting BFD2 activity had a nearly identical effect to BFD1; however, when comparing the degree of differential expression, we observed that, on average, the magnitude of changes caused by BFD1-knockdown surpassed that of BFD2 (**Fig. 1G**). These results suggest that BFD1 and BFD2 function together to remodel the transcriptome during differentiation.

### BFD2 is critical for differentiation

To rule out a dominant-negative effect from C-terminally tagging BFD2, we replaced the *BFD2* coding sequence with a TdTomato expression cassette, creating a clean knockout (Δ*bfd2*, **Fig. 2A**). Knockouts were made in a tagged BFD1-complement background (Δ*bfd1*::*BFD1-Ty*) to facilitate detection of endogenously regulated BFD1 (Waldman et al., 2020). We compared DBL staining following 48 h of alkaline stress in the parental strain, Δ*bfd1*, and Δ*bfd2*. Whereas deletion of BFD1 completely blocked formation of the bradyzoite cyst wall, close observation revealed that many Δ*bfd2* vacuoles were faintly stained with DBL (**Fig. 2B**). Measuring the intensity of DBL staining of each vacuole further emphasized the inability of Δ*bfd2* parasites to robustly upregulate this hallmark of differentiation (**Fig. 2C**).

**Figure 2.**
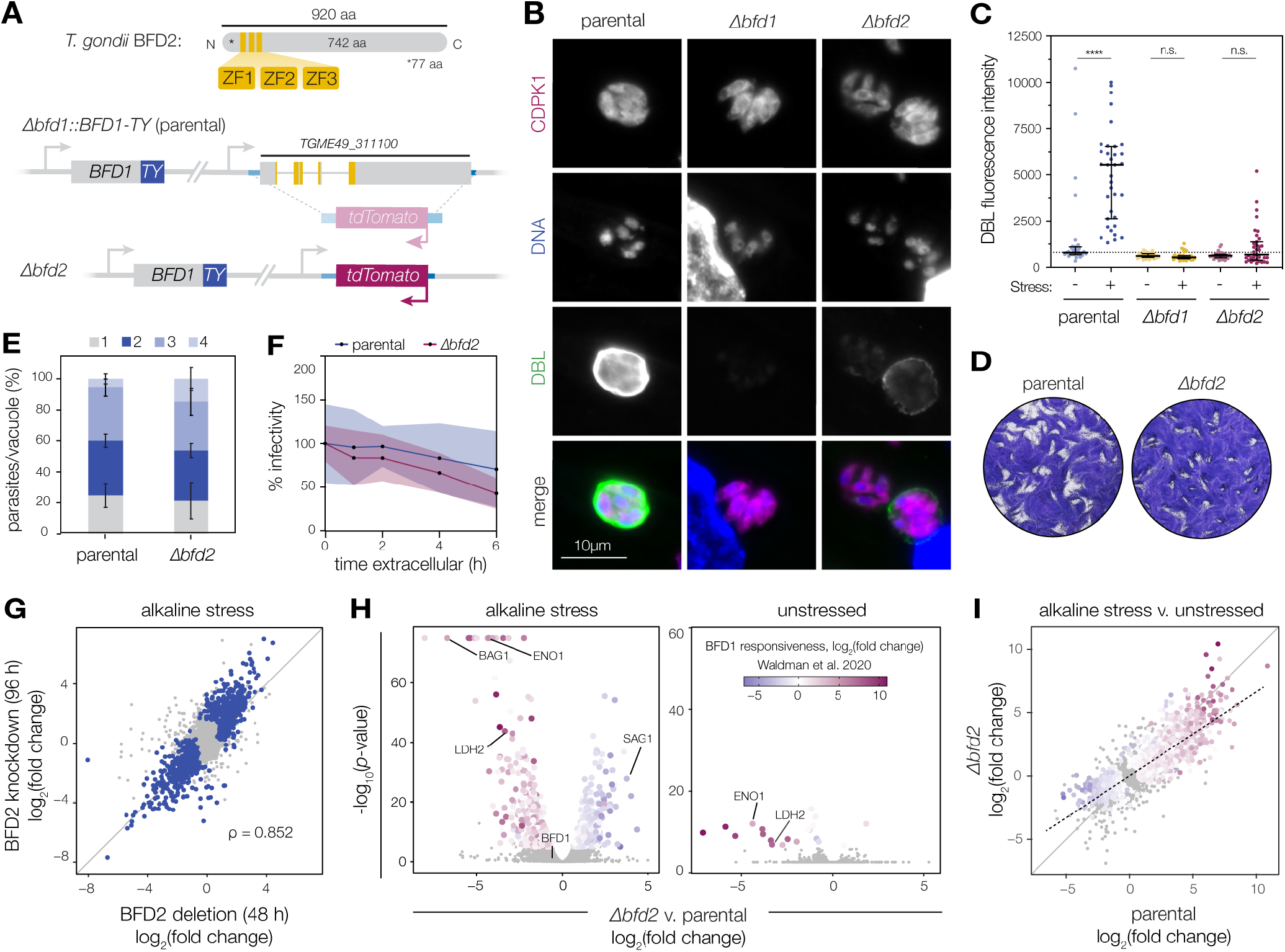
BFD2 is a CCCH-type zinc finger protein required for differentiation. (**A**) Diagram of BFD2 (top) highlighting tandem CCCH zinc finger domains (yellow). Generation of Δ*bfd2* (bottom) in a parental strain expressing TY-tagged *BFD1*. The endogenous *BFD2* coding sequence was replaced with a *TdTomato* fluorescence cassette. (**B**) Representative vacuoles after 48 h of alkaline stress. CDPK1 labels individual parasites (magenta), DNA is stained with Hoechst (blue), and differentiated vacuoles are stained with DBL (green). (**C**) DBL fluorescence intensity after 48 h under unstressed or alkaline-stressed conditions. Quantitative data reflect measurements from a single biological replicate but are representative of observations from multiple experiments. Median ± interquartile range (IQR) plotted for at least 34 vacuoles per condition. (**D**) Plaque assays of the indicated strains grown under unstressed conditions for 16 days. (**E**) Replication assays of parental and Δ*bfd2* parasites. Mean ± s.d. plotted for *n* = 3 biological replicates. The proportion of vacuoles containing 1, 2, 4, or 8 parasites is not significantly different between strains. (**F**) Parasite infectivity following varying periods of extracellular incubation. Infectivity is plotted relative to parasites directly isolated from host cells (t = 0). Mean ± s.d. plotted for *n* = 5 separate fields counted from one independent infection series. Data do not meet statistical significance at any time point. (**G-I**) Effect of BFD2 deletion on the chronic-stage transcriptome. Data are based on n = 3 independent infections. Differentially regulated genes (adjusted *p*-value < 0.05 calculated by DESeq2) are highlighted. Agreement of transcriptional changes in BFD2-knockdown parasites (analyzed in **Fig. 1**) and Δ*BFD2* after 96 h and 48 h in alkaline stress, respectively. Spearman correlation was performed on the union of differentially regulated genes in both samples (blue). (**H**) Differential expression (DE) analysis of Δ*bfd2* parasites after 48 h under alkaline-stressed or unstressed conditions. Color assigned by log2(fold change) during conditional BFD1 expression (Waldman et al., 2020). (**I**) Comparison of the effects of alkaline stress on Δ*bfd2* and its parental strain (Δ*bfd1::BFD1-Ty*). Significantly affected genes are colored as in **H**. Trend line fit by linear regression highlights the dampened transcriptional response of Δ*bfd2*. \*\*\*\**p* < 0.0001, one-way ANOVA with Tukey’s method for multiple comparisons.

In contrast to our prior observations with Δ*bfd1*, parasites lacking BFD2 displayed slightly reduced plaque formation (**Fig. 2D**). However, this represents a minor fitness defect because Δ*bfd2* replication appeared normal by other measures, including regular passaging, cell division (**Fig. 2E**), and the ability to withstand extracellular stress (**Fig. 2F**). Motivated by this observation, we revisited the results of our previous CRISPR-based screens. Although BFD2 appears dispensable under standard growth conditions in the RH strain (Sidik et al., 2016), in the ME49 screen which identified BFD1, four of the five guide RNAs targeting BFD2 dropped out over serial passaging in standard media (Waldman et al., 2020) (**Fig. S2**). The same guides were further depleted from differentiating populations; however, the mixed signal from the different guide RNAs and the apparent effect of BFD2 on ME49 fitness caused it to be excluded as a potential regulator of differentiation in our prior work.

To better understand the role of BFD2 throughout the *Toxoplasma* asexual cycle, we performed RNA sequencing on Δ*bfd2* and its parental strain under alkaline-stressed or unstressed conditions. Rank-order correlation showed strong agreement between Δ*bfd2* and the non-functional knockdown strain (*BFD2-mNG-mAID)* under alkaline stress (**Fig. 2G**). Specifically, 753 genes were dysregulated in Δ*bfd2* parasites under stress, with the majority reflecting a clear BFD1-responsive signature (**Fig. 2H-I**). By contrast, we observed only minor differences in unstressed cultures, also limited to BFD1-regulated genes like *ENO1* and *LDH2* (**Fig. 2H**). This observation is consistent with an absence of spontaneously differentiated parasites in the Δ*bfd2* population, which normally form at low frequency in wildtype ME49 strains (DBL-positive vacuoles in unstressed samples, **Fig. 2C**). The partial differentiation of Δ*bfd2* parasites was reflected in their transcriptional response to stress, which mirrored that of the parental strain, albeit with a marked decrease in magnitude (**Fig. 2I**). Together, these data validate our observations from the conditional allele, suggesting that BFD2 is required to fully establish the BFD1 transcriptional program.

### Parasites lacking BFD2 are unable to generate cysts in mice

Since the triggers of differentiation may be different in cell culture compared to vertebrate infection, we examined how BFD2 impacts virulence and cyst formation in mice. Female CD-1 mice were infected by intraperitoneal injection with 100 tachyzoites of either the parental or Δ*bfd2* strain (**Fig. 3A**). Despite the modest fitness defect of Δ*bfd2* in vitro, weight loss and mortality were equivalent between the two cohorts (**Fig. 3B-C**), indicating that BFD2 is dispensable for normal progression of acute *Toxoplasma* infection. We assessed cyst burden in the brains of all mice, based on the coincidence of DBL and a general parasite marker (CDPK1; **Fig. 3D**). After 45 days, all surviving animals injected with the parental strain harbored hundreds of cysts; however, none were detected for Δ*bfd2* (**Fig. 3E**). Histology agreed with these results (**Fig. 3F**). In moribund animals, cysts from the parental strain were first observed 23 days post-injection, but none were ever found for the mutant. This eliminates the possibility that Δ*bfd2* cysts form and are cleared during early stages of infection. Interestingly, despite the absence of cysts, localized inflammation and evidence of pathology was still detected in the brains of Δ*bfd2-*infected animals, characterized by lymphocyte infiltration, hemorrhaging, and brain calcification (**Fig. 3F**). These results could imply that low level replication of mutant parasites occurs without differentiation, triggering persistent immune activation. We conclude that BFD2 is dispensable for the acute infection in mice, but necessary for chronic differentiation.

**Figure 3.**
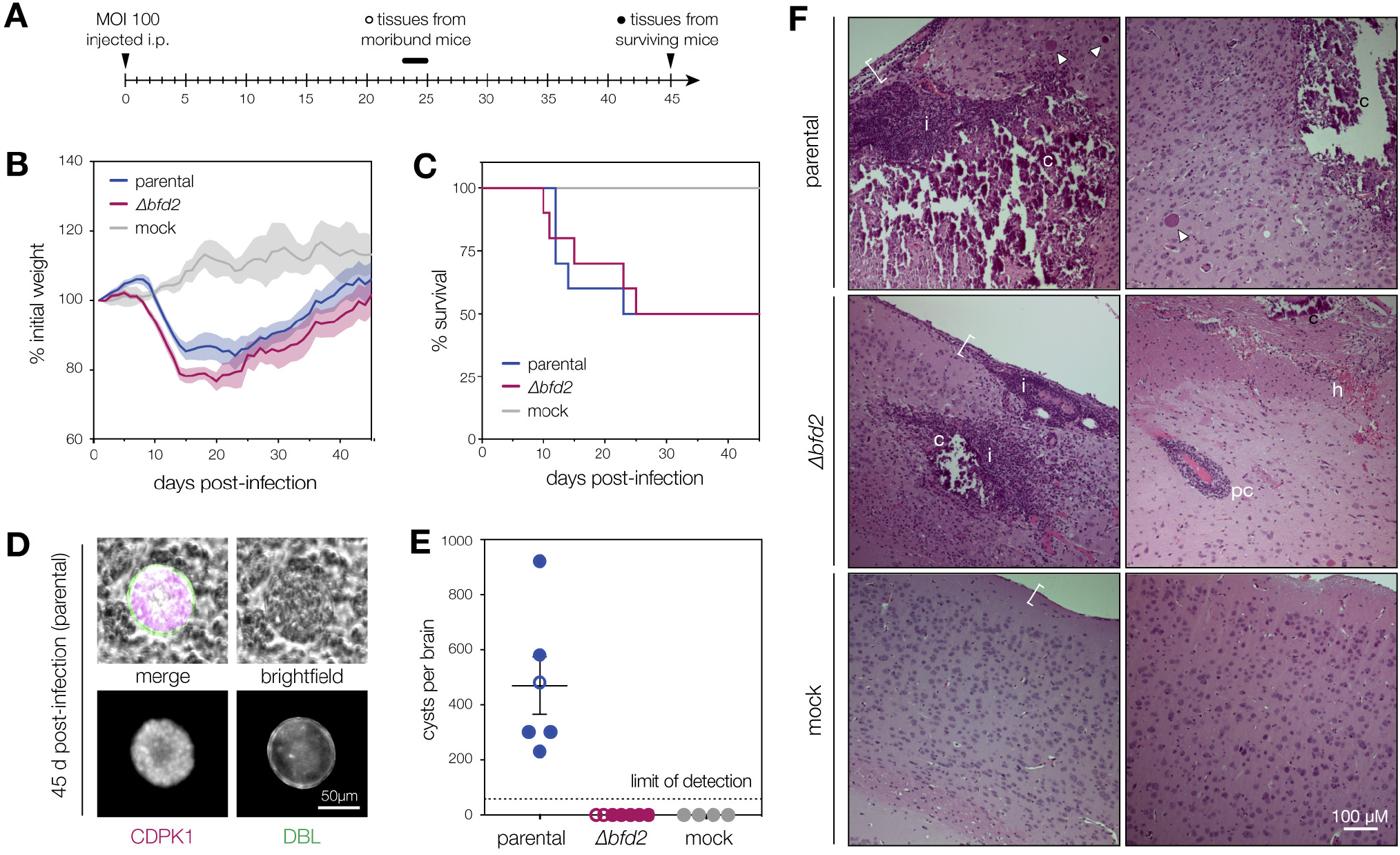
Parasites lacking BFD2 fail to produce brain cysts in mice. (**A**) Timeline of mouse infections. CD-1 female mice were injected i.p. with 100 tachyzoites from each strain (*n* = 10) or mock injected with PBS (*n* = 4). Cyst burdens were assessed in all moribund animals at the time of euthanasia and in surviving animals 45 days post-infection. (**B**) Normalized weights of animals in each infection group. Mean ± s.e.m. plotted for all surviving animals at a given time point. (**C**) Survival curve of infected animals, time-matched to weights displayed in **B**. (**D**) Representative cyst produced by the parental strain. The cyst wall is stained with DBL (green) and individual parasites are stained with CDPK1 (magenta). (**E**) Cyst burden per infected brain. Graphs reflect counts from surviving animals (closed circles) as well as all moribund animals sacrificed after 23 days from the time of infection (open circles). (**F**) Histology of brain sections from infected and mock-infected animals. Two representative images are shown per strain with signs of pathology indicated: calcification (c), immune cell infiltration (i), perivascular cuffing (pc), hemorrhaging (h), and meningeal inflammation (bracket). Cysts are indicated with white arrows.

### BFD1 and BFD2 operate as a positive-feedback loop that promotes differentiation

BFD2 is transcriptionally responsive to BFD1 and its promoter shows clear evidence of BFD1 binding— characteristics which should place it downstream in the regulatory hierarchy. However, given the analogous effects of these two factors on differentiation, we sought to test their relationship more rigorously. Stress-responsive changes in *BFD2* transcript levels were quantified by RT-qPCR in either the presence (Δ*bfd1::BFD1-Ty*; wildtype) or absence of BFD1 (Δ*bfd1*), as well as in parasites expressing a nonfunctional version lacking DNA-binding motifs (Δ*bfd1::BFD1*^ΔMYB^*-Ty*)(Waldman et al., 2020). In line with previous transcriptional profiling, *BFD2* was robustly upregulated in wildtype parasites under alkaline stress, exhibiting an approximate 6-fold increase after 2 days (**Fig. 4A**). Although both the knockout and nonfunctional-complement strain retained a low-level of induction, the magnitude of this effect was substantially diminished (∼2-fold), confirming that regulation of BFD2 is indeed mediated by BFD1. By contrast, *BFD1* levels were not significantly affected by either alkaline stress or deletion of BFD2.

**Figure 4.**
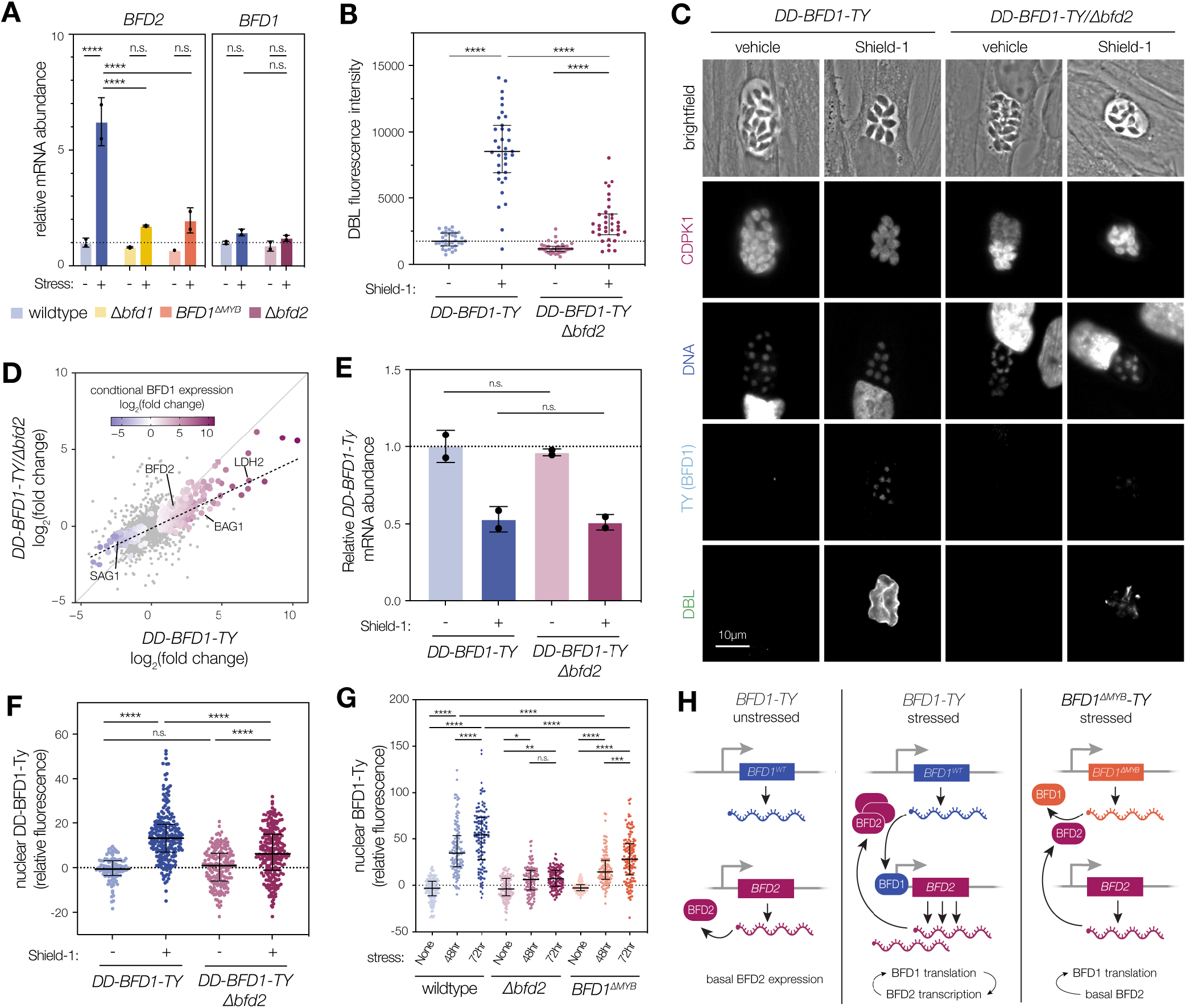
BFD1 and BFD2 comprise a positive-feedback loop. (**A**) RT-qPCR of *BFD2* and *BFD1* mRNA in parasites after 48h in the presence or absence of stress. For simplicity, strains are labeled as wildtype (Δ*bfd1::BFD1-Ty*) or *BFD1*^ΔMYB^ (Δ*bfd1::BFD1*^ΔMYB^*-Ty*) in reference to the version of BFD1 expressed. For relative quantification of either transcript, data were analyzed by comparative CT using *GCN5B* as an internal control and unstressed Δ*bfd1::BFD1-Ty* parasites as a reference sample. Complete ΔCT values and analysis are provided in **Table S4**. Mean ± s.d. plotted for *n =* 2 biological replicates. (**B**) Quantitation of DBL staining following 48 h in 3 μM Shield-1 or vehicle. Measurements were performed for a single biological replicate but are representative of observations from multiple infections. Median ± IQR are shown for at least 34 vacuoles per condition. (**C**) Representative vacuoles after 48 h in Shield-1 or vehicle. Parasites are labeled with CDPK1 and DNA stained with Hoechst. BFD1 is detected by the TY epitope. DBL specifically stains differentiated vacuoles. (**D**) Comparison of significantly regulated genes in *DD-BFD1-Ty* and *DD-BFD1-Ty/*Δ*bfd2* parasites grown for 48 h in 3 μM Shield-1 relative to vehicle alone. Genes meeting the statistical threshold in either sample (adjusted p < 0.05 calculated by DESeq2) are colored by their transcriptional responsiveness to conditional BFD1 expression (Waldman et al., 2020). Trend line fit by linear regression. (**E**) Quantification of nuclear BFD1 staining in **C**. Median ± IQR is shown for *n* = 107–226 parasite nuclei. (**F**) Relative abundance of *DD-BFD1-Ty* mRNA after 120 h in 3 μM Shield-1 or vehicle. Transcripts were quantified as in **A** using vehicle-treated *DD-BFD1-Ty* as the reference sample. Mean ± s.d. plotted for *n* = 2 biological replicates. ΔCT are listed in **Table S5**. (**G**) Quantification of nuclear BFD1 staining under alkaline-stressed or unstressed conditions. Wildtype (Δ*bfd1::BFD1-Ty*) and *BFD1*^ΔMYB^ (Δ*bfd1::BFD1*^ΔMYB^*-Ty*) labels refer to the version of BFD1 expressed. For each strain and condition, median ± IQR is plotted for *n* = 98–126 nuclei. (**H**) Model of the positive feedback loop between BFD1 and BFD2. In acute stages (left), BFD2 is expressed at a basal level. *BFD1* is transcribed but protein is not stably produced. Under alkaline stress (center), BFD2 promotes expression of BFD1, causing activation of the bradyzoite program; in turn, increased transcription of *BFD2* by BFD1 further increases the capacity for BFD1 expression. In parasites expressing a nonfunctional BFD1 (right), BFD2 still facilitates *BFD1* expression under stress but, because robust transcriptional activation of *BFD2* does not occur, BFD1 cannot be fully induced. **** p < 0.0001, *** p < 0.001, ** p < 0.01, * p < 0.05; One-way ANOVA with Tukey’s method for multiple comparisons.

To assess the stress-dependency of this relationship, we asked whether loss of BFD2 could also hamper differentiation induced by conditional BFD1 expression. Previously, we generated a regulatable version of BFD1 (*DD-BFD1-Ty*, **Fig. S3**) that is constitutively degraded unless stabilized by the small molecule Shield-1. Treatment with Shield-1 leads to accumulation of BFD1 protein, causing uniform differentiation in the absence of stress (Waldman et al., 2020). In this genetic background, *BFD2* was knocked-out and differentiation was assessed by DBL staining following 48 h under Shield-1-or vehicle treatment. As expected, culturing *DD-BFD1-Ty* in the presence of Shield-1 resulted in stage conversion, with over 94% of vacuoles showing DBL staining above vehicle-treated controls (**Fig. 4B– C**). Although we still observed a weak DBL signal in the knockout, addition of Shield-1 could not completely overcome the loss of *BFD2* (**Fig. 2C**)—reminiscent of the incomplete differentiation induced with alkaline stress in Δ*bfd2* parasites. This result was mirrored at the transcriptome level, which reflected an inability of *DD-BFD1-Ty/*Δ*bfd2* to fully induce many BFD1-regulated genes (**Fig. 4D**). Thus, BFD2 appears to be required for the BFD1 program regardless of whether it is triggered by stress or through the conditional expression of BFD1. Despite being regulated downstream of BFD1, we considered the possibility that BFD2 may be required for robust BFD1 induction. To test this, we verified that conditional expression of BFD1 was equivalent in the parental and Δ*bfd2* strains. Loss of BFD2 had no measurable effect on *DD-BFD1-Ty* mRNA abundance (**Fig. 4E**), despite a slight, Shield-1–induced decrease in transcription that could be ascribed to the heterologous promoter (alpha tubulin) driving the transgene (Waldman et al., 2020). However, immunostaining for DD-BFD1-Ty revealed a reduction in BFD1 protein levels in Δ*bfd2* parasites (**Fig. 4C**). This observation was quantified by measuring the signal intensity in individual parasite nuclei (**Fig. 4F**), suggesting that BFD2 is required to maximally induce expression of the master regulator. Given *BFD1* mRNA levels were unaffected by the presence of BFD2 (**Fig. 4A & 4E**), we conclude that BFD2 must regulate either translation or turnover of BFD1 protein.

Our data suggested that BFD1 and BFD2 reciprocally promote one another’s expression, constituting a positive feedback loop. If this is true, it follows that full induction of BFD1 should depend on its own activity. To test this model, we examined BFD1 accumulation in the endogenous context, in wildtype, Δ*bfd2*, and Δ*bfd1::BFD1*^ΔMYB^*-Ty* parasites. When both BFD1 and BFD2 were present (Δ*bfd1::BFD1-Ty*; wildtype), BFD1 was robustly detected in alkaline-stressed parasites after 48 h, and continued to accumulate over time. Parasites lacking BFD2 also showed a small but significant increase in BFD1 protein under stress; however, the absolute level was dramatically reduced, reaching only ∼7% of that measured for the wildtype strain after 72 h (**Fig. 4G**). This was consistent with the weak induction of chronic-stage genes we observed by transcriptional profiling of Δ*bfd2* (**Fig. 2H-I**). Intriguingly, the nonfunctional complement strain, Δ*bfd1::BFD1*^ΔMYB^*-Ty*—which harbors basal levels of BFD2, that do not substantially change in response to stress— accumulated an intermediate amount of BFD1. Thus, we conclude that basal levels of BFD2 facilitate stress-responsive expression of BFD1. This in turn promotes transcription of BFD2, closing the positive feedback loop (**Fig. 4H**).

### BFD2 directly mediates translation of *BFD1* mRNA

The cytosolic localization we observed for BFD2-mNG-mAID (**Fig. 1C**) is consistent with its proposed role in post-transcriptional regulation. However, given that BFD2 activity was disrupted by C-terminal tagging (**Fig. 1D**), we sought to verify this localization with a functional transgene. We complemented *Δbfd2* by introducing an N-terminally tagged cDNA copy into the endogenous locus (*HA-BFD2;* **Fig. 5A**). *HA-BFD2* parasites displayed normal plaque formation (**Fig. 5B**) and complete rescue of DBL staining under alkaline stress (**Fig. 5C-D**), indicative of successful complementation. As was observed for the C-terminally tagged allele, HA-BFD2 was detectable by immunofluorescence in acute-stage parasites and upregulated during differentiation (**Fig. 5C**). Additionally, HA-BFD2 localized to the cytosol irrespective of culture conditions, suggesting that its activity is likely confined to the cytosol.

**Figure 5.**
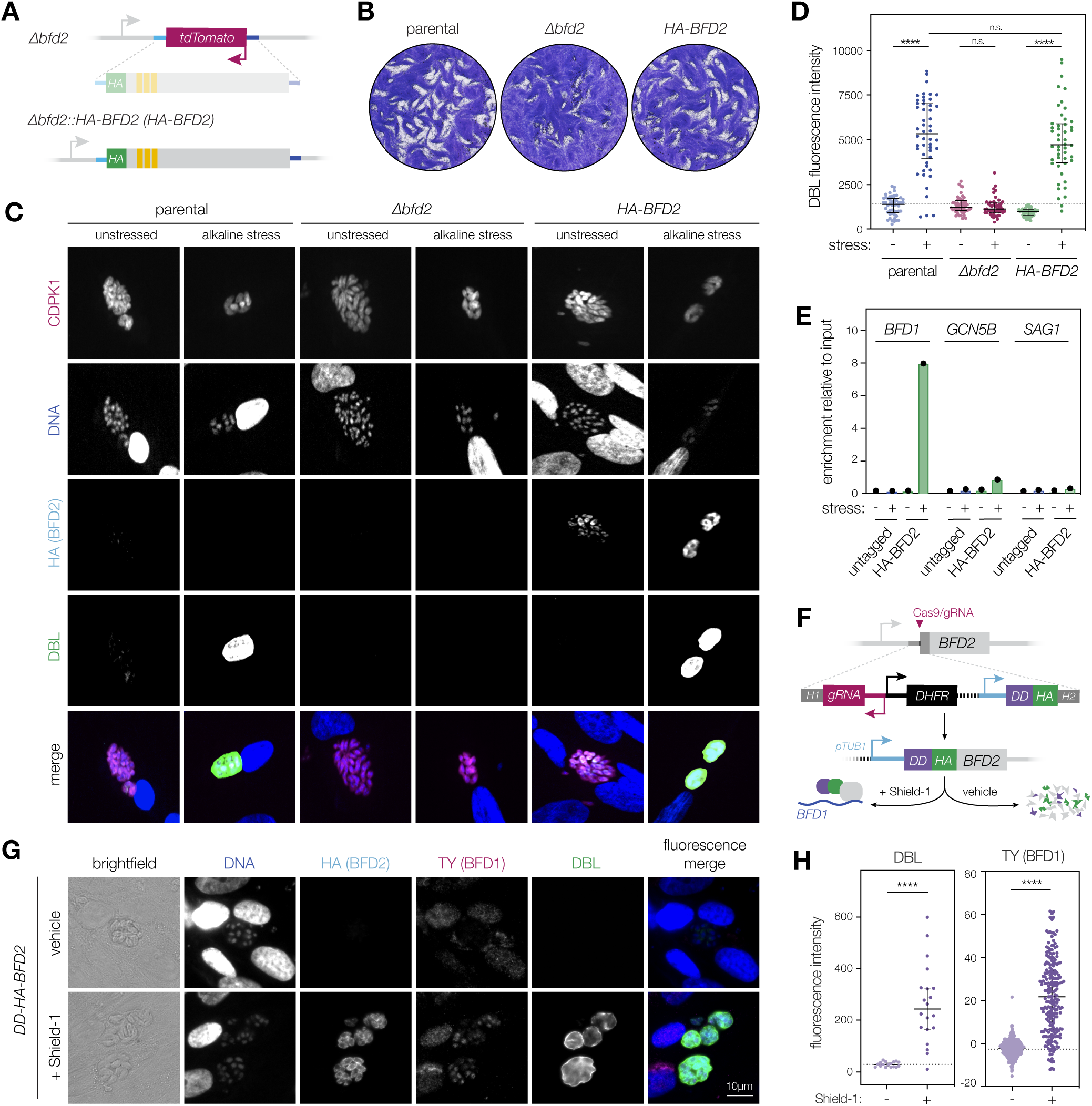
BFD2 interacts with the *BFD1* transcript and is sufficient for BFD1 translation. (**A**) Complementation of Δ*BFD2*. An HA-tagged cDNA copy of *BFD2* was knocked-in at the endogenous locus, replacing the *TdTomato* cassette. (**B**) Plaque assays of the indicated strains grown under unstressed conditions for 16 days. Parental refers to the Δ*bfd1::BFD1-Ty* strain, used to construct Δ*bfd2*. (**C**) Representative vacuoles after 48 h of alkaline stress. Individual parasites are labeled with CDPK1 (magenta) and DNA with Hoechst (blue). BFD2 is detected by the HA epitope (cyan). Differentiated vacuoles are stained with DBL (green). (**D**) Quantitation of DBL fluorescence after 48 h under alkaline-stressed or unstressed conditions. For each strain and condition, data reflect the median ± IQR calculated *n* = 46–50 vacuoles. (**E**) Analysis of BFD2-bound transcripts after 72 h under alkaline-stressed or unstressed conditions. Pull-downs of HA-BFD2 were performed using antibodies specific to the HA epitope, and the indicated transcripts were quantified by RT-qPCR with normalization by the Percent Input method. For ΔCT values and analysis, refer to **Table S6**. (**F**) Generation of a conditionally stabilized BFD2 overexpression strain. The alpha tubulin promoter (*pTUB1*) drives transgene expression while an N-terminal destabilization domain (DD) enables regulatable induction of the gene product. Accumulated BFD2 is detected by the HA epitope. Additional strain details are provided in Materials & Methods and **Fig. S4**. (**G**) Representative vacuoles after 48 h of treatment with 3 μM Shield-1 or vehicle. DNA is stained with Hoechst (blue), BFD2 is detected by the HA epitope (cyan), and BFD1 by TY (magenta). Differentiated vacuoles are stained with DBL (green). (**H**) Analysis of DBL staining and BFD1 protein accumulation in the samples shown in **G**, with median ± IQR are plotted. In vehicle and Shield-1-treated samples respectively, *n* = 17 and 20 vacuoles (DBL) or *n* = 236 and 216 individual nuclei (BFD1). Quantification was performed for a single experiment, but data are representative of observations from multiple independent infections. **** p < 0.0001; one-way ANOVA with Tukey’s method for multiple comparisons.

Given the putative function of BFD2 as an RNA-binding protein, we considered that BFD2 might interact directly with the BFD1 transcript. To test this, we performed immunoprecipitation using antibodies recognizing the HA epitope, which should specifically enrich for HA-BFD2 but not an untagged allele. To distinguish between constitutive and stress-dependent interactions, we harvested cells cultured under each condition. Total RNA was extracted from enriched and unenriched input samples, and analyzed by RT-qPCR for *BFD1* and two negative controls—*GCN5B*, a gene whose expression is not affected by alkaline stress, and *SAG1*, an acute-stage gene downregulated during differentiation. Strikingly, although all transcripts were detected in the input, only *BFD1* mRNA was enriched in the HA-BFD2 pulldown (**Fig. 5E**). Moreover, the association was detected specifically under stress. Since *BFD2* deletion prevents accumulation of BFD1 protein, but has no obvious impact on *BFD1* mRNA abundance (**Fig. 4D-G**), we posit that BFD2 facilitates the translation of *BFD1* through a stress-specific association with the transcript.

### Overexpression of BFD2 triggers differentiation

In keeping with our model of BFD2 as a positive regulator of BFD1 translation, we considered that conditional overexpression of BFD2 could drive differentiation. We generated a regulatable version of BFD2 through modification of the native locus, replacing the endogenous promoter with that of alpha tubulin while appending an N-terminal destabilization domain to the coding sequence (*DD-HA-BFD2*; **Fig. 5F**). To facilitate detection of BFD1, constructs were introduced into the Δ*bfd1::BFD1*^WT^*-TY* parental strain. Treatment with the stabilizing Shield-1 ligand led to robust cytosolic accumulation of DD-HA-BFD2 (**Fig. 5G**) and by 48 h, 95% of vacuoles developed DBL positivity above the level of the vehicle-treated control (**Fig. 5H**). Importantly, BFD1-TY also accumulated in the nuclei of parasites treated with Shield-1 (**Fig. 5G & H**), further establishing BFD2 as a driver of BFD1 protein expression. Thus, we posit that overexpression of BFD2 is indeed sufficient for differentiation; however, the effect is likely a reflection of the hierarchical relationship between the two factors, with BFD2 activating translation of BFD1 which, in turn, orchestrates the bradyzoite transcriptional program.

## DISCUSSION

The formation of bradyzoite cysts in an infected host is essential for chronic persistence of *Toxoplasma*. We recently identified BFD1 as the master regulator of this process (Waldman et al., 2020). Now we expand upon this model with the discovery of BFD2, a second factor indispensible for the chronic stage. BFD2 is a cytosolic RNA-binding protein that is expressed in acute stages and upregulated during differentiation. We show that basal levels of BFD2 are required for induction of BFD1, placing it upstream in the regulatory hierarchy. Conversely, BFD1 promotes transcription of *BFD2*, constituting a positive feedback loop. We show that BFD1 protein levels—but not mRNA abundance— are affected by BFD2 knockout, implying translational regulation. In agreement with this, BFD2 interacts with the *BFD1* transcript under alkaline stress. Taken together, our data support a model wherein BFD2 licenses stress-dependent translation of *BFD1* and acts in a positive feedback loop with the master regulator to drive chronic differentiation.

Several studies have described RNA- and DNA-binding proteins that influence rates of stage conversion (Gissot et al., 2013, 2017; Hong et al., 2017; Huang et al., 2017; Liu et al., 2014; Radke et al., 2013, 2018; Vanchinathan et al., 2005; Walker et al., 2013); however, their relationship to the master regulator of differentiation, BFD1, has not been established. Our own studies identified several direct targets of BFD1 with nucleic acid-binding domains (Waldman et al., 2020), suggesting a hierarchy of differentiation-promoting factors. It was therefore unexpected that, of the five candidates examined in this work, few exhibited any obvious transcriptional signature. AP2IX-9 was previously shown to inhibit the expression of bradyzoite-specific mRNAs *BAG1, LDH2*, and *ENO1* (Hong et al., 2017; Radke et al., 2013); yet, we observed no consequence to its depletion under alkaline stress. This may be explained by the observation that AP2IX-9 is only transiently expressed during the earliest stages of differentiation (Radke et al., 2013). Similarly, although prior work implicated *AP2IB-1* as transcriptionally upregulated in chronic forms, our inability to detect the tagged protein suggests that it may not be translated at the time points sampled. Profiling mutants outside the 96 h differentiation window used in our study could have captured effects for these factors in the developing bradyzoite population. Ironically, the candidate with the most obvious transcriptional signature was BFD2, which appears to function as a regulator of translation. In this case, disruption led to widespread dysregulation of chronic-stage mRNAs, as a consequence of its relationship to BFD1. Much subtler changes were detected for the other RNA-binding proteins, although our results do suggest that TGME49_224630 may be associated with alkaline stress responses outside the BFD1-regulated differentiation program. In light of our observations from BFD2, we suspect that examining the impact of these RNA-binding factors at the proteome level may reveal additional layers of regulation.

BFD2 belongs to a widespread class of RNA-binding proteins defined by the presence of one or more CX7-8CX4-5CX3H (CCCH) zinc finger motifs (Hall, 2005). These domains are found in proteins with diverse functions, affecting virtually all stages of RNA metabolism (Fu and Blackshear, 2017; Hajikhezri et al., 2020; Maeda and Akira, 2017). However, few CCCH proteins have been functionally characterized in apicomplexans. Two recent studies in *Plasmodium* spp. implicated ZFP4 and Pb103 in sexual reproduction due to gametocyte-specific effects (Hanhsen et al., 2022; Hirai et al., 2021). Loss of ZFP4 prevented exflagellation and led to a downregulation of many male gametocyte-specific mRNAs. Pb103 itself was shown to be translationally repressed during female gametogenesis; yet the mechanism of regulation conferred by either factor has not been definitively shown. Intriguingly, a CCCH protein in the non-apicomplexan parasite *Trypanosoma brucei*, known as TbZFP3, was found to promote differentiation through regulated association with polyribosomes (Paterou et al., 2006; Walrad et al., 2009)—consistent with positive translational control such as we have described for BFD2. A domain search of the *T. gondii* genome reveals at least 33 other genes encoding putative CCCH motifs (VEuPathDB.org). Given their importance across diverse parasite phyla, we suspect that further exploration of this protein family will expose new factors with key roles in stage conversion.

Our data suggest that BFD2 is a core part of the chronic differentiation program in *Toxoplasma*. While orthologs of BFD2 can be found in other closely related cyst-forming apicomplexans, such as *Neospora* and *Hammondia*, it does not appear to be conserved outside of the *Sarcocystadae* family (VEuPathDB.org). BFD1 displays a similar pattern of conservation, suggesting that the relationship between the two factors may be maintained in closely related species. Whereas loss of BFD1 completely blocks differentiation, vacuoles containing Δ*bfd2* parasites still exhibit low-level DBL staining under alkaline stress, consistent with the formation of incipient cysts in cell culture. This is further supported by transcriptional profiling, which reflects weak induction of the bradyzoite program. However, no cysts—immature or otherwise—were ever identified in mice infected with Δ*bfd2*, either by analysis of brain homogenates or by histology. This disparity likely reflects differences between natural differentiation and in vitro culture systems, suggesting that the conditions are harsher for cysts in the context of vertebrate infection.

BFD2 is expressed in both proliferative and latent stages of *T. gondii*, implying additional roles outside of differentiation. Although survival of *BFD2* knockouts demonstrates that the factor is not essential for tachyzoite growth, we did observe a modest fitness cost in culture in the ME49 strain. The limited differences in the transcriptomes of unstressed Δ*bfd2* and wildtype parasites could be attributed to the lack of spontaneous differentiation in the former. Replication rates, sensitivity to extracellular stress, and acute-stage virulence were also comparable, leaving the deficiency of Δ*bfd2* unknown. Notably, whereas loss of *BFD2* in our prior ME49 screens led to similarly compromised fitness (Waldman et al., 2020), deletion in the RH (type I) lineage of *Toxoplasma* appears innocuous (Sidik et al., 2016). This suggests that strain-specific differences might account for the defects of Δ*bfd2* tachyzoites. Additionally, based on the demonstrated function of BFD2 as a translational regulator, we suspect that its effects on other genes, besides *BFD1*, are likely born out at the proteome (rather than transcriptome) level. A more complete characterization of BFD2 clients could reveal its function in acute stages.

We show that BFD2 interacts with the *BFD1* transcript in a stress-dependent manner, suggesting that it functions as a positive regulator to license *BFD1* translation. This could potentially be mediated through interaction with translation initiation factors or ribosomal recruitment— possibilities we are continuing to explore. While the mechanisms controlling differential binding of BFD2 to its mRNA targets will require further investigation, we note that conditional overexpression of BFD2 is sufficient to drive differentiation irrespective of stress. This relationship does not preclude the function of BFD1 as the master regulator of the chronic stage, since our collective data suggest that differentiation depends on the transcriptional changes induced by BFD1. However, it does indicate that the regulatory machinery involved is responsive to the concentration of BFD2 and can be overwhelmed stoichiometrically. Based on this observation, we propose that once the amount of BFD2 exceeds a certain threshold (e.g., via transcriptional activation by BFD1), maintenance of the bradyzoite state becomes less dependent on the signals that initiated it. This phenomenon—known as hysteresis—is well-described in other developmental programs, such as cell-type determination in multicellular organisms and commitment to cell division (Ferrell, 2002; Lek et al., 2010; Moses and Rubin, 1991; Sha et al., 2003; Solomon, 2003). In the context of the *T. gondii* chronic stage, such a circuit could provide a stabilizing force to maintain parasites in a differentiated state in the presence of variable stimuli.

Feedback loops endow genetic circuits with several notable features. The discovery of a positive feedback loop acting on BFD1 therefore has fundamental implications for the properties of *T. gondii* latency. In addition to achieving signal amplification, these circuits frequently exhibit bistability. That is, whereas hysteresis renders a genetic system less susceptible to noise, bistability confers switch-like, binary control (Ferrell, 2002). Previously, it was suggested that acute-to-chronic stage conversion in *T. gondii* involves the production of intermediate slow-growing tachyzoite stages, termed pre-bradyzoites (Hong et al., 2017; Radke et al., 2013, 2018). This notion is not inconsistent with a binary model, since bistable systems can encode more than two cell states due to non-equilibrium (Fang et al., 2018). Nevertheless, our observation that Δ*bfd2* parasites partially induce the chronic-stage program but cannot persist as cysts in mice suggests that robust binary commitment is important. Perhaps more intriguingly, bistable systems exhibiting hysteresis are capable of ‘remembering’ a stimulus after it has been removed. Indeed, network theory suggests that a double-positive loop such as the one we describe for BFD1 and BFD2 should lock into a self-perpetuating steady-state (Ferrell, 2002). This raises the question of how the bradyzoite program is overturned during reactivation. Although the conditions leading to reactivation during natural infection are not well understood, alkaline-stressed parasites returned to standard media will resume proliferation. We speculate that this may require additional reactivation factors which antagonize the activity of BFD1 or BFD2. Alternatively, it is possible that the absence of stress may be sensed directly by the circuit. Other groups have proposed that differentiation requires a balance of regulators, and identified proteins that repress the transcription of bradyzoite-specific genes (Hong et al., 2017; Radke et al., 2013, 2018). However, whether these or other factors directly inhibit BFD1 or BFD2 has yet to be examined.

In summary, our work shows that BFD2 is a major determinant of *T. gondii* persistence due its involvement in a positive feedback loop with the master regulator, BFD1. This discovery opens up exciting new questions surrounding the regulation of the BFD1 program, particularly as it relates to the maintenance and reactivation of chronic stages. Further exploration of the BFD1-BFD2 circuit will likely uncover heuristics that govern timing and commitment in differentiation. Together, these advances will aid in the design of therapeutic interventions to prevent or reverse chronic stages, bringing us closer to a radical cure against toxoplasmosis.

## MATERIALS & METHODS

### Parasite and host cell culture

*T. gondii* parasites were maintained in human foreskin fibroblasts (HFFs) grown at 37°C under 5% CO2 in standard medium, consisting of DMEM (GIBCO) supplemented with 3% heat-inactivated fetal bovine serum (IFS), 2 mM L-glutamine (GIBCO), and 10 μg/ mL gentamicin (Thermo Fisher). Routine passaging of ME49 parasites was performed on a 2-3 day cycle— prior to lysis of host cell monolayers—by scraping and transfer to fresh HFFs. For all experimental infections, parasites were released from host cells after scraping by extrusion through a 27-gauge needle (referred to as mechanical disruption), and inoculated onto HFFs in standard medium supplemented with 10% IFS. For differentiation, parasites were switched to alkaline-stress medium (RPMI 1640 supplemented with 1% IFS, 10 μg/mL gentamicin, 50 mM HEPES, and pH-adjusted to 8.1 with 10 N NaOH) and maintained under ambient CO2. Cultures were provided fresh medium every 2 days in all experiments lasting longer than 48 h. IAA was used at 50 μM in standard media and 500 μM under alkaline stress. Shield-1 was used at 3 μM in standard medium (Banaszynski et al., 2006).

### *T. gondii* transfection

Parasites were mechanically released from host cells using a 27-gauge needle and passed through a Millex filter unit (5 μm pore size, Merck Millipore) to remove host cell debris. Parasites were pelleted by centrifugation at 1000 x *g* for 10 minutes, resuspended in cytomix electroporation buffer (10mM KPO4, 120 mM KCl, 150 mM CaCl, 5 mM MgCl, 25 mM HEPES, 2mM EDTA) supplemented with 2 mM ATP and 5 mM glutathione, and combined with the constructs to be transfected in a final volume of 400 μL. Electroporation was performed in a 4mm cuvette using an ECM 830 Square Wave electroporator (BTX) with the following settings: 2 pulses, 1.7 kV, 176 μs pulse length, and 100 ms interval.

### *T. gondii* strain generation

Oligos were ordered from IDT. A full list can be found in **Table S2**. All cloning was performed with Q5 Master Mix (NEB) and NEBuilder HiFi DNA Assembly Master Mix. Genotyping PCRs were performed using Taq DNA polymerase in standard buffer.

#### TIR1-expressing ME49 (ME49/TIR1)

Starting with the *ME49*Δ*ku80*Δ*hxgprt* strain (Waldman et al., 2020), we introduced the heterologous TIR1 auxin receptor into a defined, neutral locus on chromosome VI (Markus et al., 2019). To facilitate integration, we used a previously validated gRNA/Cas9-expression plasmid targeting the neutral locus (Genbank #MN019116). A repair template encoding TIR1 under the control of the alpha tubulin promoter (*pTUB1*) and a CAT expression cassette was amplified from Addgene plasmid #87258 using oligos P1/P2, which include overhangs homologous to the regions immediately adjacent to the gRNA target site. Parasites co-transfected with the gRNA and repair construct were selected in chloramphenicol-containing media and, after 2 weeks, surviving parasites were sub-cloned by dilution plating in 96-well plates. Wells containing single plaques were screened for successful integration events by PCR using P3/P4.

#### TGME49_200385, TGME49_311100, TGME49_306620, TGME49_253790, TGME49_224630, and TGME49_208020 mNG-mAID-tagged strains

In *ME49/TIR1*, individual genes were endogenously tagged with *mNG-mAID* using the previously described high-throughput (HiT) tagging strategy (Smith et al., 2021). Cutting units specific to each candidate were purchased as IDT gBlocks (P5–P10) and assembled with the empty pGL015 mNG-mAID HiT vector backbone (GenBank: OM640005, **Fig. S4A**).

Approximately 75 μg of each assembled vector was BsaI-linearized and co-transfected with 50 μg of the pSS014 Cas9-expression plasmid (GenBank: OM640002). Bulk populations of each strain were selected for 1 week in standard medium containing pyrimethamine, and parasites were subcloned into 96-well plates. Single-plaque-containing wells were screened for successful integration at the respective endogenous locus using a common reverse primer in the mNG tag (P11), and a gene-specific forward primer (P12–17) binding upstream of the integration site. PCR products were sequenced to verify maintenance of the reading frame across the junction where recombination occurred.

#### BFD2-knockout strains (Δbfd2 and DD-BFD1-TY/ Δbfd2)

Oligos encoding two different gRNAs against the 5′ (P18/P19) and 3′ (P20/P21) ends of the endogenous *BFD2* coding sequence were annealed and assembled with the pU6-Universal Cas9-expression plasmid (Genbank #OM640003), and sequenced with P22. A repair template consisting of *pTUB1* driving expression of TdTomato was amplified using primers P23/P24, which encode 40 bp of homology to regions immediately up- and downstream of the gRNA target sites. 25 μg each of the repair template and both gRNA/ Cas9-expression plasmids were transfected into the *ME49*Δ*Ku80*Δ*bfd1::BFD1-TY* and *ME49*Δ*Ku80*Δ*bfd1/ HXGPRT::pTUB1-DD-BFD1-TY* strains (previously described in Waldman et al., 2020) to generate Δbfd2 and *DD-BFD1-TY/Δbfd2*, respectively. Transfectants were maintained for 2–3 days to allow for construct integration and expression of the reporter, then *BFD2*-knockouts were FACS-sorted from the bulk population directly into 96-well plates based on fluorescent signal. Single-plaque wells were screened for both the presence of the integrated TdTomato cassette (P25/ P26) and the absence of the endogenous *BFD2* coding sequence (P25/P27).

#### ME49ΔKu80Δbfd1::BFD1-TYΔbfd2::HA-BFD2 (HA-BFD2)

*HA-BFD2* was reintroduced into the endogenous locus such that complemented parasites could be isolated based on loss of TdTomato using FACS. Two gRNAs were designed against the 5′ and 3′ ends of the TdTomato cassette, and ordered as oligos (P28/P29 and P30/P31, respectively) for annealing and Gibson-assembly with the pU6-Universal plasmid. For a repair template, *BFD2* was amplified from cDNA (P32/P33) and assembled with PCR fragments spanning ∼500 nt immediately up-(P34/P35) and downstream (P36/ P37) of the *BFD2* coding sequence. The N-terminal HA epitope was introduced in-frame at the junction between the upstream amplicon and *BFD2*, through overhangs designed into the corresponding reverse and forward primers (i.e., P35 and P32). The annealed product was then used as a template to generate ∼15 μg of linear repair construct (PCR-amplified with P38/P39), which was co-transfected with 20 μg of each gRNA plasmid into Δbfd2 parasites. After an initial round of FACS to enrich for low-or non-fluorescent cells, complemented mutants were sorted directly into 96-well plates for subcloning. Single-plaque wells were screened by PCR for reintroduction of the tagged version of *BFD2* (P25/P27) and deletion of the TdTomato cassette (P25/ P26). Positive mutants were additionally verified by sequencing through the junctions where recombination occurred, ensuring reconstitution of the endogenous sequences flanking *HA-BFD2*.

#### Conditional overexpression of BFD2 (DD-HA-BFD2)

The conditional BFD2-overexpression strain was generated using a derivation of the HiT vector strategy described previously (Smith et al., 2021), modified for manipulation of the N-(rather than C-) termini of targeted genes. A cutting unit specific to *BFD2* was purchased as an IDT gBlock (P40) and Gibson-assembled with the empty pHL053 pTUB1-DD HiT vector backbone (GenBank accession number pending, **Fig. S4B**). The resultant plasmid was sequence-verified with P41, then BsaI-linearized and transfected into *ME49ΔKu80Δbfd1::BFD1-TY* parasites along with 50 μg the pSS014 Cas9-expression plasmid. Mutants were selected for 1 week in standard medium containing pyrimethamine prior to subcloning by dilution plating into 96-well plates. Clonal populations were PCR-screened using primers to both detect successful integration of the construct (P42/P27) and verify absence of the wildtype allele (P43/P27).

### Immunofluorescence assays

Prior to inoculation with *Toxoplasma*, HFFs were seeded onto coverslips and grown in standard media until confluent (∼2-3 days). At the time of fixation, samples were washed twice with PBS to remove extracellular parasites, then fixed with 4% formaldehyde for 20 min, permeabilized with 1% Triton X-100 for 8 min, and blocked for 25 min with a solution composed of 5% IFS and 5% normal goat serum in PBS (blocking buffer). Primary and secondary antibody incubations were performed for 1 h in a humidified chamber, with coverslips inverted on 30 μL of antibody diluted in blocking buffer and spotted onto Parafilm. All steps from fixation through staining were performed at room temperature, with three PBS washes between. After the final wash, coverslips were rinsed in deionized water and mounted onto slides with 5 μL of Prolong Diamond (Thermo Fisher). The mountant was cured for 45 min at 37°C or overnight at room temperature before imaging. All antibody and stain dilutions were prepared as follows: rabbit anti-GAP45 (gift from Dominique Soldati, University of Geneva), 1:10,000; guinea pig anti-CDPK1, custom antibody (Corvance), 1:10,000; mouse anti-mNeonGreen, clone 32F6 (ChromoTek), 1:500; mouse anti-Ty (Bastin et al., 1996), 1:1000; rabbit anti-HA, clone 71-5500 (Invitrogen), 1:1000); mouse anti-HA, clone 16B12 (Biolegend), 1:1000; Hoechst 33258 (Santa Cruz), 1:20,000; DBL-488 (Vector Labs), 1:1000. Secondary antibodies labeled with Alexa Fluor 488, 594, or 647 (Thermo Fisher) were all used at 1:1000. Images were acquired using an Eclipse Ti (Nikon) microscope or an RPI spinning disc confocal microscope equipped with NIS Elements and Metamorph software, respectively.

### Plate-based differentiation screen

Parasites were inoculated onto HFFs in black-wall 96-well microplates and allowed to invade for 4 h in standard media. Plates were washed twice with PBS and switched to alkaline-stress medium containing 500 μL IAA or an equivalent volume of PBS (vehicle). After 48 h, plates were fixed and permeabilized using the conditions described above. Without blocking, wells were then treated sequentially with 50 μL of primary (guinea pig anti-CDPK1) and secondary (anti-guinea pig-594, DBL-488, and Hoechst 33258) staining solutions for 1 h each, with three PBS washes after all steps. Fluorescence images were collected at 20x magnification with a Cytation3 Cell Imaging Multi-Mode Plate Reader (Aligent Techonologies). Scans were focused on the nuclear stain using laser auto focus, with subsequent channels imaged on a 3 μm offset.

Differentiation rates were scored blindly using a custom-built automated image analysis pipeline. Briefly, a mask was created for each set of images using the CDPK1 counterstain to delineate individual vacuoles, regardless of differentiation status. This was applied to the corresponding DBL scan from each set, creating a second mask to enable visualization of the isolated DBL-staining profile for each vacuole. Using the Interactive Learning and Segmentation Toolkit (ilastik v. 1.3.3)(Berg et al., 2019), vacuoles were then categorized as negative-intermediate-or high, based on the intensity and variance of the DBL signal. Data represent the mean and standard deviation of three independent experiments. The pipeline was built and trained on hand-scored differentiation assays prior to its application in this study.

### RNA-seq analysis of conditional knockdown strains

HFFs in 15 cm dishes were inoculated with syringe-released parasites under standard conditions. After 4 h, the monolayer was rinsed with PBS and parasites were switched to alkaline-stress media containing either 500 μM IAA or an equivalent volume of PBS (vehicle). Cultures were harvested at 96 h, with fresh medium provided after 2 days. Parasites were mechanically released by sequential extrusion through a 27- and 30-gauge needle, then filtered through a 5 μm filter to remove host cell debris. Cell suspensions were pelleted by centrifugation at 1,000 x *g* for 10 min and resuspended in Trizol before flash-freezing in liquid nitrogen for short-term storage. Total cellular RNA was isolated using the Zymo Direct-zol RNA Miniprep Kit and RNA quality was assessed using a BioAnalyzer or Fragment analyzer. For all samples, cell pellets were split in two at the time of RNA isolation and treated as technical replicates in downstream processing. Libraries were generated using the SWIFT RNA Library Kit and sequenced on one lane of the NovaSeqS4 with 150×150 paired-end reads. Reads were trimmed to remove adapter sequences using Cutadapt v.3.6 and aligned to the ME49 genome (ToxoDB v.56 assembly) with STAR v.2.7.0a. Differential expression analysis was done using the DESeq2 package (v. 1.12.3) in R. Adjusted *p-value* of 0.05 was used as a statistical cutoff for differential expression.

### RNA-seq analysis of BFD2 deletion

HFFs in 15 cm dishes were inoculated with parasites in standard medium. After 4 h, the monolayer was rinsed with PBS and cultures were either returned to standard medium for acute stages or treated with one of two differentiation triggers. For Δ*bfd1::BFD1-Ty* and Δ*bfd1::BFD1-TyΔBFD2* (**Fig. 2E-F**), stage-conversion was induced by alkaline stress. For *DD-BFD1-Ty* and *DD-BFD1-Ty/ΔBFD2* (**Fig. 4D**), parasites received standard medium supplemented with either 3 μM Shield-1 to induce conditional BFD1 expression of the equivalent volume of 100% ethanol (vehicle). Parasites were grown for an additional 48 h then mechanically released from host cells by scraping and serially passing through 27- and 30-gauge needles, followed by filtration with a 5 μm filter. Cell suspensions were pelleted by centrifugation at 1,000 x *g* for 10 min and flash-frozen in liquid nitrogen. Total cellular RNA was isolated using the Qiagen RNeasy Plus Micro Kit following the protocol for animal and human cells. For all samples, cell pellets were split into two at the time of RNA isolation and treated as technical replicates in downstream processing. Library preparation, sequencing, and analysis were performed with the same pipeline as was applied to conditional knockdown strains (see above).

### Extracellular stress assays

Single-cell suspensions of extracellular parasites were generated by syringe-release from host cells with a 27-gauge needle followed by filtering through a 5 μm filter. 50 μL of the suspension was inoculated onto coverslips seeded with HFFs in standard media, and the remaining volume was maintained at 37°C. At 1, 2, 4, and 6 h timepoints following the initial infection, additional coverslips were inoculated with the parasite suspension at an equivalent MOI. All coverslips were fixed with formaldehyde after 24 h of growth, stained for DNA (Hoechst 332580) and CDPK1 as described in *Immunofluorescence assays*, and multiple fields of view were imaged for each coverslip using an Eclipse Ti microscope. The infectivity of parasites at each timepoint was quantified as a measure of the number of vacuoles containing 2 or more parasites per field, normalized to the number of host cell nuclei per field. For either strain, data were normalized to the infectivity of freshly-released parasites (relative to t = 0).

### Intracellular replication assays

HFFs on coverslips were inoculated with parasites for 4 h, then washed twice with PBS and returned to standard media. After 24 h from the time of infection, samples were fixed, permeabilized, and stained for DNA (Hoechst 33258) and CDPK1 as described under *Immunofluorescence assays*. For each sample, the number of parasites per vacuole was calculated from a minimum of 100 vacuoles. Data represent the mean of three independent infections.

### Plaque assays

HFFs in 6-well plates were inoculated with 5×10^3^ parasites and allowed to sit undisturbed in standard media supplemented with 10% IFS for 16 days. Plates were washed twice with PBS and fixed at room temperature for 10 min with 100% ethanol. Plates were then dried overnight and the wells stained for 5 min with 2 mL of crystal violet solution (12.5 g crystal violet, 125 mL 100% ethanol, 500 mL 1% ammonium oxalate), followed by two washes with water.

### Quantitative analysis of immunofluorescence assays

Coverslips were prepared as described under “Immunofluorescence assays” and imaged at 60x magnification. All analyses were performed using Fiji (Schindelin et al., 2012). For DBL quantification, individual vacuoles were manually defined as regions of interest (ROIs) based on a CDPK1 counterstain. ROIs were saved with the Fiji ROI Manager function, applied to the corresponding DBL channel, and their cumulative pixel intensity (Integrated Density) was quantified. Pixel intensity was also quantified over non-infected regions of the monolayer, providing a measure of Mean Background Intensity for each image. The Corrected Total Fluorescence (CTF) for individual ROIs was then calculated using the following function: CTF = [Integrated Density – (ROI area × Mean Background Fluorescence)]/ROI area. Nuclear BFD1 staining was quantified similarly, with parasite nuclei manually delineated as ROIs based on Hoechst staining. Mean Background Intensity was measured from non-nuclear regions of intravacuolar parasites.

### Assessment of acute pathogenicity and brain cyst formation

#### Mice

Six week-old female CD-1 mice were purchased from Charles River and maintained in the Whitehead Institute for three weeks prior to their use in experiments. All animal research facilities were accredited by the Association for Assessment and Accreditation of Laboratory Animal Care and all protocols were approved by the Institutional Animal Care and Use Committee at the Massachusetts Institute of Technology.

#### Mouse infections

Mice were infected intraperitoneally by injecting 100 µL of PBS containing 100 acute-stage parasites of either the parental strain (Δ*bfd1::BFD1-Ty*), Δ*bfd2*, or mock (PBS alone). Weights were recorded daily as a measure of disease severity. Animals displaying excessive distress or morbidity were euthanized with CO2 followed by cervical dislocation. Surviving animals were sacrificed 45 days post-infection; *n* = 10 for groups infected with each parasite strain and *n* = 4 for mock-infected animals.

Brains were harvested from all mice at the time of euthanasia. Brains from moribund animals were used in their entirety for cyst quantification. For animals surviving to day 45, brains were bisected at the time of harvest and one hemisphere was set aside for examination by immunohistochemistry (see below).

#### Ex vivo quantification of bradyzoite cysts

Brain tissues were dissected into 2 mL PBS and homogenized by repeated extrusion through an 18-gauge needle until a uniform consistency was achieved (approximately 10x). Homogenates were brought to 10 mL with PBS and pelleted by centrifugation at 1,000 x *g* for 5 min. The supernatant was discarded, homogenates were resuspended in 700 µL PBS and the final volume was recorded. 100 µL of homogenate was then combined with 900 µL of ice-cold 100% methanol to fix for 5 min at room temperature. The samples were centrifuged at 5,200 x *g* for 5 min (conditions used for all subsequent centrifugation steps), washed with 1 mL PBS, centrifuged again, and the supernatant was removed by aspiration. To label cysts, the pelleted samples were resuspended in 500 uL of primary staining solution (DBL-488 and guinea pig anti-CDPK1, diluted 1:150 and 1:1000 in PBS, respectively) and incubated overnight at 4°C, with rotation. The following day, samples were centrifuged, washed with 1 mL PBS, and incubated in secondary staining solution (anti-guinea pig-594, diluted 1:1000 in PBS) rotating at room temperature. After 1 h, the homogenates were washed again and resuspended in 1 mL PBS. For each sample, four 50 µL aliquots were plated into a clear-bottomed 96-well microplate and examined at 20x magnification by fluorescence microscopy. Cysts were identified on the basis of their double-positivity for both DBL and CDPK1. With the exception of those prepared from moribund mice, all samples were quantified blindly. Cyst burdens represent the mean of the four counts collected for each sample, multiplied by the appropriate dilution factor.

#### Immunohistochemistry

Brain tissue was fixed in 10% formalin, embedded in paraffin and sectioned. Deparaffinized sections were stained for hematoxylin and eosin according to standard protocol for evaluation of tissue pathology. Images were acquired at 10x magnification with a Leica DM6000 Widefield Fluorescence Microscope.

### Quantitative reverse transcription PCR (RT-qPCR)

Parasites were inoculated into T-12.5 flasks seeded with HFFs and allowed to invade for 4 h, then switched to standard or alkaline-stress medium. At the time of harvest (48 h post-infection for unstressed cultures; 48 h or 120 h for alkaline-stressed), monolayers were rinsed twice with PBS to remove extracellular parasites and the entire contents of each flask were collected. Samples were centrifuged at 1,000 x *g* for 10 min, the supernatant was discarded, and pellets were flash-frozen in liquid nitrogen. Total cellular RNA was isolated from frozen pellets using the Qiagen RNeasy Plus Micro Kit following the protocol for animal and human cells. To eliminate contaminating genomic DNA, RNA was subjected to DNaseI digestion using the TURBO DNA-free kit (Thermo Fisher). RNA samples were then quantified by NanoDrop spectrophotometer and used for first strand cDNA synthesis performed with SuperScriptIII (Thermo Fisher) and random hexamer priming. Dye-based qPCR was performed using PowerUp SYBR Green Master Mix (Thermo Fisher) and included a ‘No-RT’ control for each sample. Reactions were run on a QuantStudio6 Real-Time PCR system (Applied Biosystems) using the “Standard” ramp speed with the following thermocycling parameters: 2 min at 50°C; 2 min at 95°C; 40 cycles of 15 s at 95°C, followed by 1 min at 60°C. All primer pairs (**Table S3**) were previously validated for amplification efficiency in the 95-105% range and their specificity verified by melt curve analysis. Transcripts of interest were quantified by comparative CT (ΔΔCT) method, using *GCN5B* as an internal control gene and normalizing to unstressed samples.

### BFD2 immunoprecipitation and analysis of enriched transcripts

*HA-BFD2* and Δ*bfd1::BFD1-TY* (untagged control) parasites were allowed to invade HFFs for 4 h under standard conditions, then switched to standard or alkaline-stress medium. At 48 and 72 h respectively, unstressed and stressed samples were rinsed with PBS and mechanically released from host cells by serially passing through 27- and 30-gauge needles, followed by filtration with 5 μm filter units to eliminate host cell debris. For each strain and condition, ∼2×10^7^ parasites were collected by centrifugation (1,200 x *g* for 10 min) and pull-downs were performed using the Magna RIP RNA Binding Protein Immunoprecipitation Kit (Sigma-Aldrich). Unenriched ‘input’ samples were generated by reserving back 10% of the starting lysate; the remaining volume was subjected to immunoprecipitation on protein A/G magnetic beads coated with 5 µg of mouse anti-HA IgG (Biolegend, clone 16B12) as described in the Magna RIP technical manual. RNA was isolated in parallel from both enriched and input samples by conventional phenol-chloroform extraction and used for first strand cDNA synthesis performed with SuperScriptIII and random hexamer priming. Dye-based qPCR was then performed as described above using 5 ng of diluted cDNA with the primers listed in **Table S3** and including No-RT controls. Enrichment of transcripts after BFD2-pull-down was calculated by the Percent of Input method.

### Statistical analysis

Information about biological replicates, the number of observations, and exact statistical tests performed can be found in figure legends. For RNA-seq analysis, adjusted *p-values* were calculated by DESeq2. For all other quantitative analyses, statistical tests were performed in Prism (GraphPad).

## ACKNOWLEDGEMENTS

We thank Dominique Soldati for the GAP45 antibody; the Whitehead Institute Genome Technology Core, particularly Tom Volkert, Jennifer Love, and Sumeet Gupta for their expertise with library preparation and next-generation sequencing; and the Whitehead Bioinformatics and Research Computing Core for consultation and software support. This work relied on VEupathDB, and we thank all contributors to this resource. This project was supported by grants from the NIH (R01AI158501) and the Smith Family Foundation (Odyssey Award) to S.L..

## AUTHOR CONTRIBUTION

This project was conceptualized by M.H.L., B.S.W., and S.L.. All experiments were performed by M.H.L., C.J.G., and L.S. Resources were provided by S.L. and C.A.H. Data analysis was performed by M.H.L., S.C., and S.L. The manuscript was prepared by M.H.L. and S.L. with input from all contributing authors.

## Notes

### Competing Interest Statement

The authors have declared no competing interest.

