## Supplementary Information for "A positive feedback loop controls *Toxoplasma* chronic differentiation"

### SUPPLEMENTAL FIGURES

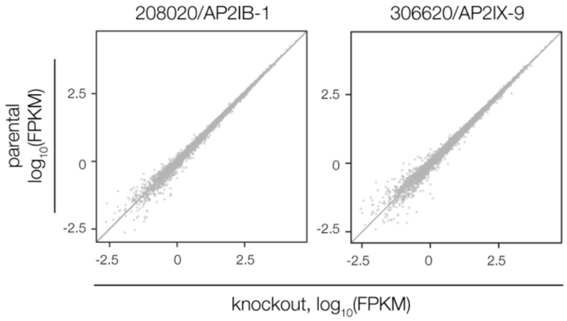


**Figure S1. Effects of AP2IX-9 and AP2IB-1 knockdown on the chronic-stage transcriptome.** Data reflect changes in knockdown strains (relative to the parental) after 96 h in alkaline-stress medium containing IAA. Differential expression analysis was performed as in **Fig. 1E**, based on $n = 2$ biological replicates. No genes are significantly affected (adjusted $p < 0.05$, calculated by DESeq2) by depletion of either factor.

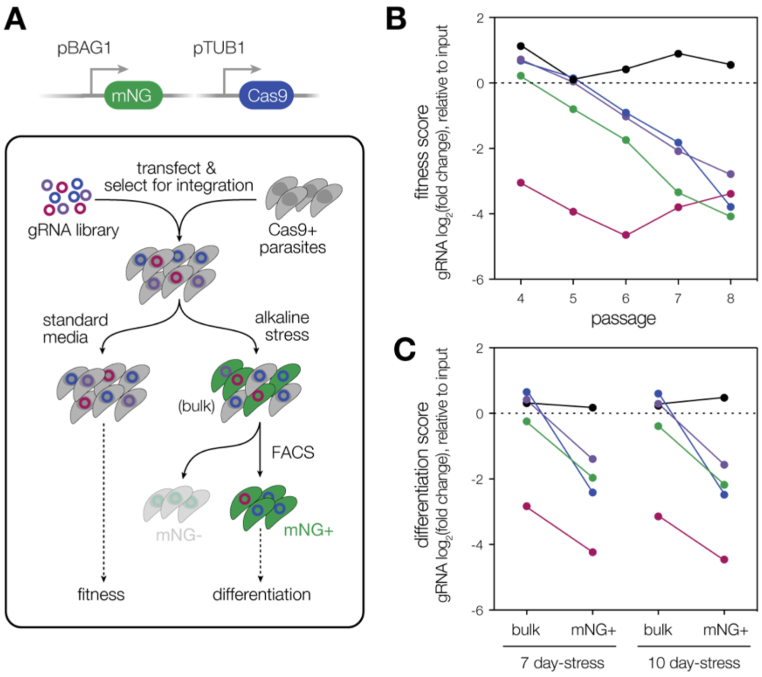


**Figure S2. Reexamining the performance of BFD2-deficient parasites in previous ME49 screens.** (**A**) Overview of the CRISPR-based screen that identified BFD1 (Waldman et al., 2020). A CRISPR-compatible ME49 strain was modified to express mNeonGreen (mNG) under the bradyzoite-specific *BAG1* promoter (pBAG1), enabling isolation of chronic stages by fluorescence-activated cell sorting (FACS; top). The reporter strain was transfected with guide RNA (gRNA) libraries targeting ~200 predicted nucleic acid-binding proteins, with 5 gRNAs per gene. After several days to allow for guide integration and inactivation of targeted genes, transfectants were split between alkaline-stressed and unstressed (standard media) conditions (bottom). Samples were collected from each population over a 10-day period, with bradyzoites ($mNG^+$-stressed parasites) isolated by FACS. Integrated gRNAs from all samples were enumerated by next-generation sequencing and the abundance of each guide was assessed relative to the input library. The $log_2$(fold change) for guides targeting each gene is referred to as its fitness or differentiation score, based on representation in unstressed or bradyzoite samples, respectively. (**B**) Analysis of BFD2-targeting gRNAs. 4 of the 5 guides targeting *BFD2* were lost from the transfectant pool under standard conditions over the course of serial passaging. Sequence-level analysis revealed that the single guide that remains abundant (black) is non-cutting. (**C**) Analysis of BFD2 gRNAs during differentiation. In samples harvested at both 7 and 10 days post induction of alkaline stress—with the exception of the non-cutting guide (black)—gRNAs targeting BFD2 are underrepresented in bradyzoite samples relative to the unsorted alkaline-stressed population (bulk).

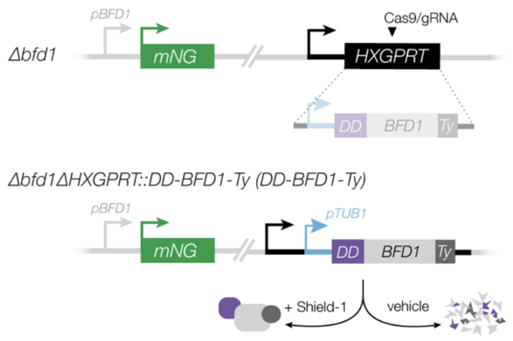


**Figure S3. Schematic of the conditional BFD1 expression strain.** *DD-BFD1-Ty* was constructed as described previously (Waldman et al., 2020), by integration into the *HXGPRT* locus in the Δ*bfd1* background. A heterologous promoter (*pTUB1*) drives expression of the transgene, but DD-BFD1-Ty protein is only stabilized upon treatment with Shield-1.

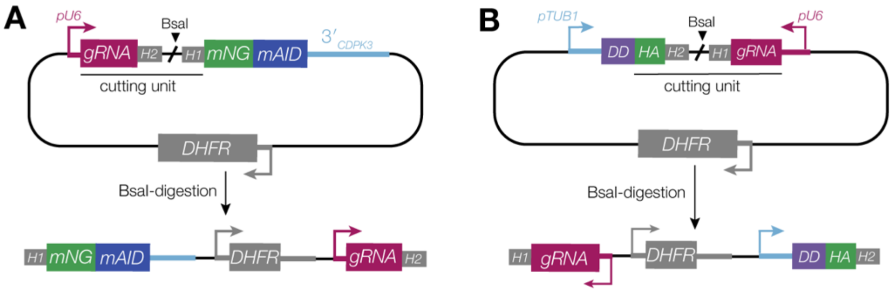


**Figure S4. Strain construction using the HiT vector strategy.** (**A**) Schematic of C-terminal tagging HiT vectors (top), previously described in Smith et al., 2021. Targeted integration of BsaI-linearized constructs (bottom) is facilitated by a guide RNA (gRNA) specific to the 3′ end of the coding sequence and 40 bp homology regions (H1, H2), both encoded in the gene-specific cutting unit. Transcription of the gRNA is driven by a type III promoter (pU6). A heterologous 3′ untranslated region ($3'_{CDPK3}$) allows expression of the gene product. *DHFR* denotes a pyrimethamine resistance cassette to enable mutant selection. (**B**) N-terminal HiT vector configuration for generation of conditional overexpression strains. Construct integration endogenously tags the targeted gene with the Shield-1-stabilized degradation domain (DD) and replaces the native promoter with that of alpha tubulin (*pTUB1*). Cutting units are designed similarly to those in **A**, with a gRNA that targets the coding sequence 5′ end encoded in reverse orientation. For the inducible BFD2 strain in particular, HA was also designed into the cutting unit to enable detection of the protein by the same epitope used for examination of endogenously regulated BFD2.

### SUPPLEMENTAL TABLES

**Table S1.** Subsets of genes significantly affected by conditional depletion of *BFD1, BFD2, TGME49_253790,* and *TGME49_224630* under alkaline stress. Related to Figure 1. The behavior of listed genes in prior transcriptional and chromatin profiling (columns labeled as ".2020") as well as all other transcriptional datasets included in this work is shown.

**Table S2.** Primers used for molecular cloning and genotyping.

**Table S3.** RT-qPCR primers used for analysis of *BFD1, BFD2, GCN5B,* and *SAG1* transcripts. Related to Figures 4 and 5.

**Table S4.** Analysis of *BFD1* and *BFD2* mRNA abundance in Δ*bfd1::BFD1*$^{\Delta MYB}$*-Ty,* Δ*bfd1::BFD1-Ty,* Δ*bfd1,* Δ*bfd1::BFD1*$^{\Delta MYB}$*-Ty,* and Δ*bfd2* parasites after 48 h under alkaline-stressed or unstressed conditions. Related to Figure 4A. Mean CT values are the average of three technical replicates.

**Table S5.** Analysis of *DD-BFD1-Ty* mRNA abundance in *DD-BFD1-Ty* and *DD-BFD1-Ty/*Δ*bfd2* parasites following 48 h of treatment with Shield-1 or vehicle. Related to Figure 4E. Mean CT values are the average of three technical replicates.

**Table S6.** Analysis of *BFD1, GCN5B,* and *SAG1* mRNA abundance in input and enriched pull-down samples. Related to Figure 5E. Data were analyzed by the Percent of Input method. Mean CT values are the average of three technical replicates.

**Table S7.** Compiled results of all differential expression analyses performed in this work. Related to Figures 1, 2 and 4.
